# Ecology of domestic dogs *Canis familiaris* as an emerging reservoir of Guinea worm *Dracunculus medinensis* infection

**DOI:** 10.1101/2020.02.19.939827

**Authors:** Robbie A. McDonald, Jared K. Wilson-Aggarwal, George J.F. Swan, Cecily E.D. Goodwin, Tchonfienet Moundai, Dieudonné Sankara, Gautam Biswas, James A. Zingeser

**Author notes:** Email addresses: Robbie McDonald, Jared Wilson-Aggarwal, George Swan, Cecily Goodwin, Tchonfienet Moundai, Dieudonné Sankara, Gautam Biswas, James Zingeser. **Correspondence author:** Robbie McDonald, Environment and Sustainability Institute, University of Exeter, Penryn Campus, Penryn, UK. **Statement of authorship:** RM, JZ, GB and DS conceived the study. RM, JW-A, GS, TM and JZ conducted the fieldwork. JW-A, CG and GS analysed samples and data. RM wrote the manuscript. JW-A, CG, GS, DS and JZ revised the manuscript. **Data accessibility statement:** Data are available at Dryad data repository https://doi.org/10.5061/dryad.vx0k6djnh.

## Abstract

Global eradication of human Guinea worm disease (dracunculiasis) has been set back by the emergence of infections in animals, particularly domestic dogs *Canis familiaris*. The ecology and epidemiology of this reservoir is unknown. We tracked dogs using GPS, inferred diets using stable isotope analysis and analysed correlates of infection in Chad, where numbers of Guinea worm infections are greatest. Dogs had small ranges that varied markedly among villages. Diets consisted largely of human staples and human faeces. A minority of ponds, mostly <200 m from dog-owning households, accounted for most dog exposure to potentially unsafe water. The risk of a dog having had Guinea worm was reduced in dogs living in households providing water for animals but increased with increasing fish consumption by dogs. Provision of safe water might reduce dog exposure to unsafe water, while prioritisation of proactive temephos (Abate) application to the small number of ponds to which dogs have most access is recommended. Fish might have an additional role as transport hosts for Guinea worm, by concentrating copepods infected with worm larvae.

**Author summary:** Guinea worm is a parasite that causes profoundly debilitating disease in humans. An eradication program has been successful in nearly eliminating the disease from people. However, the same worm has now been found in domestic dogs and the frequency of detecting Guinea worm in dogs has been increasing. This means that to eradicate Guinea worm, the infection must be eliminated in dogs as well as in people. However, not much is known about the disease in dogs. This study is the first to investigate dog ecology in relation to Guinea worm infection. We worked in the worst-affected country, Chad. We attached GPS collars to dogs to track their ranging and use of water bodies and analysed their diets using a forensic technique, based on analysing stable isotope compostion of their whiskers and potential food items. We showed that dogs living in households that provided water to their animals had a lower risk of having had Guinea worm and that dogs that ate more fish had an increased risk. These findings suggest there is a classical route for worm transmission in dogs, via drinking contaminated water, as well as a novel route, potentially by eating fish carrying a source of infection.

## Introduction

Guinea worm *Dracunculus medinensis* is a nematode parasite that causes Guinea worm disease (dracunculiasis) in humans. It was once widespread in Asia and Africa [1] but an eradication campaign has made impressive progress in reducing human cases from 3.5 million per annum in 21 countries in 1986, to only 28 in 2018, in Chad (17), South Sudan (10), and Angola (1) [2]. Alongside the near-absence of human cases, however, infections were detected in 1040 domestic dogs *Canis familiaris* and 25 domestic cats *Felis catus* in Chad, 11 dogs, five cats and one olive baboon *Papio anubis* in Ethiopia, and 18 dogs and two cats in Mali in 2018 [2] Adult Guinea worms emerging from humans and these non-human animals are genetically indistinguishable [3].

Prior to re-emergence in 2010, no human cases of Guinea worm were reported in Chad for ten years [4], suggesting an unknown reservoir of infection in non-human animals and/or undetected infection in humans, though surveillance of the disease in humans was also problematic at that time. Given the current high number of dogs infected relative to human cases and improved human surveillance in Chad, it is clear that non-human animals, particularly dogs, now constitute a contemporary reservoir of infection. Little is known about the epidemiology of Guinea worm in its non-human hosts. Although direct transmission between dogs and humans is not possible, dogs clearly maintain a persistent source of potential infection for humans in their shared environments. Thus, to eradicate Guinea worm, transmission of infection must be interrupted in non-human as well as human hosts [5]. Alongside civil insecurity in the remaining endemic areas, non-human animal infections are now the major impediment to Guinea worm eradication [6].

The pathways by which dogs acquire infection remain unknown, though two main possibilities exist. First, similar to the classical route for humans, dogs may acquire infection from drinking water containing infected copepods as the intermediate hosts. Second, dogs might consume a paratenic or transport host, most likely a fish or amphibian, that has itself eaten infected copepods [7]. In Chad, dogs drink from permanent and transient surface water bodies that often contain large copepod populations. Large-scale human exploitation of fish and frogs also provides the potential for dog consumption of their uncooked remains [7].

Options for interrupting dog infections are broadly similar to those for humans, and include: detection and containment of hosts, treating water bodies with the organophosphate temephos (Abate), and limiting exposure by reducing consumption of contaminated water and, potentially, food [8]. Since different approaches are required to implement such interventions when aimed at free-ranging dogs, it is important that risk factors are identified, to help inform management. To understand the epidemiology and control of Guinea worm in dogs, we undertook a study of their ecology in Chad, to characterise their ranging and access to water, as well as their diets and consumption of aquatic foods. We show that both a classical route of transmission, relating to exposure to potentially unsafe water, as well as a novel route, via consumption of fish, potentially as a paratenic host, influence the risk of dog infection and highlight management implications for proactive water treatment.

## Material and methods

### Ethics statement

This study was approved by the University of Exeter College of Life and Environmental Sciences (Penryn Campus) Ethics Committee (2016/1488). The Committee and the project adhered to the “Guidelines for the treatment of animals in behavioural research and teaching” of the Association for the Study of Animal Behaviour.

### Study area and subject recruitment

Fieldwork was conducted between 24 June and 12 July 2016 in three settlements (Kakale, Magrao and Largana), each comprised of multiple villages, situated in Guelendeng sous-préfecture, close to the Chari River, Chad (Figure 1). Settlements were selected because of the high numbers of infections in dogs and relative ease of access. The chief of each village provided approval and dog owners individually provided consent. For each household, we recorded the GPS location and conducted a questionnaire, recording whether dogs were taken hunting (reported by owners as entering the bush to capture wild game, and classified as hunting or not) and whether water was provided for animals within the household. For each dog, we recorded sex, age in months, body condition (on a scale of 1 = emaciated to 9 = severely obese [9] and then classified as poor ≤2 or moderate ≥3), and whether the dog had history of emergent adult Guinea worms. The Chad Guinea Worm Eradication Programme (CGWEP) provided additional surveillance data identifying dogs with confirmed infections.

**Figure 1:**
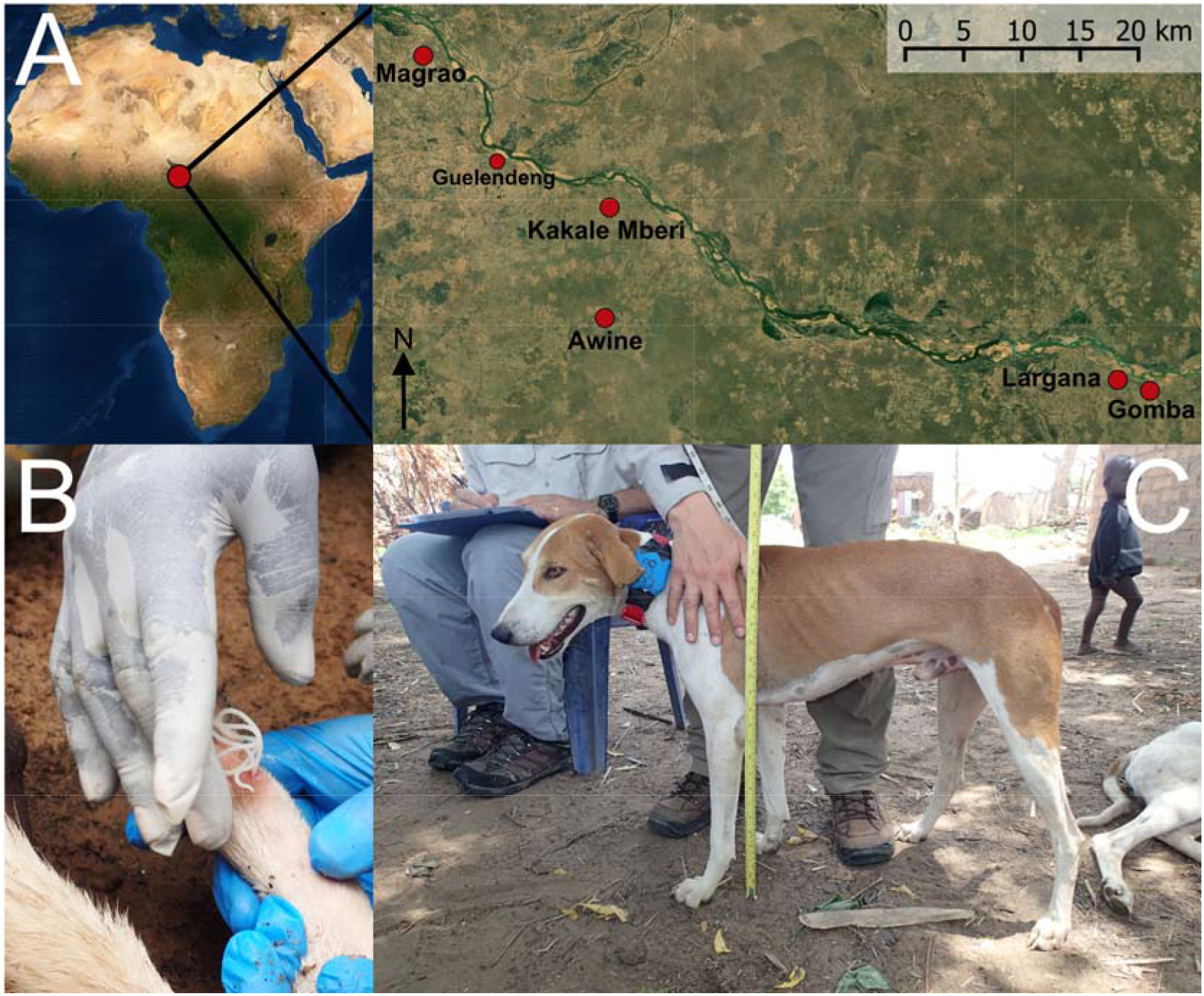
A) Locations of dog study sites on the Chari River, Chad. B) Adult female Guinea worm *Dracunculus medinensis* being extracted from a domestic dog. C) Typical dog fitted with GPS tracking collar. The settlements were: Magrao (10°59’44.31”N, 15°29’29.27”E) which encompasses the associated permanent village of Sawata, Largana (10°45’15.26”N, 16° 1’39.86”E) and the associated permanent village of Gomba (10°44’46.22”N, 16° 2’46.70”E), and Kakale-Mberi (10°53’0.79”N, 15°38’8.45”E) and the associated seasonal settlement of Awine (10°48’6.34”N, 15°37’56.61”E). Guelendeng is the major town in the sous-préfecture. Photographs Jared Wilson-Aggarwal. The satellite image was generated using the Esri world imagery basemap (sources: Esri, DigitalGlobe, GeoEye, i-cubed, USDA FSA, USGS, AEX, Getmapping, Aerogrid, IGN, IGP, swisstopo, and the GIS User Community).

### Dog tracking

All dogs, apart from pups that were <6 months old and were too small to be collared, were fitted with adjustable nylon collars (Ancol, Walsall, UK) equipped with an iGotU GT600 GPS logger (Mobile Action Technology, Taipei) with a 10 minute fix interval. After deployment for up to 14 days, collars were retrieved and data downloaded. Locations before 6am the day after deployment and after 6am the day before retrieval were excluded to avoid interference arising from our activities. Fixes that suggested speeds greater than 20km/hr were considered spurious and excluded. In one village (Gomba) in Largana, owners removed collars early, so locations after midnight prior to their removal of collars were excluded. The dogs’ core and total ranges were characterised by 60% kernel density estimates and 100% minimum convex polygons, respectively.

DigitalGlobe satellite imagery was obtained for February, October and December 2012 for Magrao, Kakale and Largana, respectively. This imagery was the nearest available in time to when our fieldwork was conducted at the end of the dry season, and likely reflects the availability of surface water earlier in the dry season. We sampled the distribution of ponds by drawing a box around all dog locations for each settlement and visually searching images for ponds. For each dog, we quantified exposure to potential sources of infection using four variables: 1. The number of GPS fixes in the vicinity (locations within 100m) of a water body (ponds and the Chari River); 2. the number of independent visits to the vicinity of water bodies (defined by an interval of 30 minutes between fixes); 3. The number of water bodies in core and total ranges; and 4. the number of different water bodies visited. For dogs with small ranges and/or those living in households close to ponds or to the Chari River, this would lead inevitably to relatively high measures of exposure, since these individuals would, in effect, be constantly ‘visiting’ the vicinity of water bodies. We considered, nevertheless, that this was an appropriate measure of variation among dogs in their relative exposure to potentially unsafe water sources. For each pond, we measured the number of dog visits, the number of dogs that visited and the number of visits by each dog (defined by an interval of 30 minutes between fixes). To characterise the ponds most visited by dogs, ponds were ranked by number of visits by all dogs and the distance of each pond to the nearest household with tracked dogs.

Variation in dog range sizes was analysed using linear models (LMs). Ranges were ln-transformed. Models contained settlement, sex, age, body condition, use in hunting activity and household water provision as explanatory variables, and the number of tracking days as a fixed effect to control for effort. Variation in the time dogs spent in the vicinity of water bodies (the number of fixes within 100 m) was analysed using a general linear model (GLM) with the same explanatory variables, except for use in hunting activity, plus the log of the number of tracking days as an offset to control for effort. A generalised additive model (GAM) related the cumulative total of dog visits per water body to the log-transformed distance from a household with tracked dogs.

### Dog diets

Composition of dog diets was determined using stable isotope analysis of dog whiskers and putative food items. Two whiskers were plucked from dogs in Kakale and Magrao during collar retrieval. The period over which whiskers had grown, and diet could be inferred, was calculated by dividing whisker length by growth rate; a subsample of dogs were fed ~50 g baits with 0.8 g (40-80 mg/kg) of rhodamine B, a food colourant that stained whiskers with a fluorescent band. Growth rate (mm/day) was calculated by dividing the distance between the whisker base and the distal end of the band by the interval between bait consumption and plucking.

To identify putative food items, owners were asked what they fed their dog, what their dog ate yesterday and what they saw other people’s dogs eating. The latter question was to identify ‘taboo’ items. Foods reported with >2% frequency were sampled, within households where possible. Otherwise, livestock meats were sampled at a market in Guelendeng, wild animal meats were obtained from hunters, and fish were sampled from fishers on the Chari. Samples were dried on the day of collection and stored in ambient conditions. Items were heat sterilised before and after importation under licence to the UK.

For each dog, one whisker was rinsed in distilled water, scraped to remove surface contaminants, and dried for 24 hours. Whiskers were cut into ~0.7 mg sections and each was weighed and sealed in a tin cup for analysis. Food samples were freeze-dried, homogenised and ~0.7 mg was weighed and sealed in a tin cup. Stable isotope analyses of carbon (δ^13^C) and nitrogen (δ^15^N) were conducted using a standard protocol [10] on a Sercon 2020 elemental analyser isotope ratio mass spectrometer. We applied a lipid-normalisation model to δ^13^C values of samples with high lipid content [11,12].

We estimated the relative contributions of putative food sources to dog diets using the SIMMR package (v.3.0) [13]. Local food source values were used where available. Isotope ratios were averaged for each dog. Trophic discrimination factors for dogs for δ^15^N (2.75‰ SD 1.19) and δ^13^C (1.64‰ SD 1.43) were obtained using the SIDER package [14]. Concentration dependence values (mean N/C) were added to the model as dogs have omnivorous diets with high variation in elemental composition among sources [15].

### Guinea worm infection

To explore correlates of infection, we analysed variation in a) whether a dog had ever been reported to have emergent Guinea worms or not, using a GLM with a binomial error structure, and b) the number of worms that had emerged from infected dogs, using a GLM with a negative binomial error structure. Explanatory variables were settlement, age, sex, body condition, household water provision and the proportion of fish in the diet estimated from stable isotope analysis. The number of GPS fixes recorded in the vicinity of water bodies, number of different water bodies visited and total range sizes were also included in model a). To maximise sample size, a staged analysis was conducted; the model was run first using data for individuals with no missing data and then run again having removed the variables that appeared in fewer than 50% of the top models, allowing the inclusion of further individuals with some missing data. The retention of diet data meant these analyses were limited to Magrao and Kakale. Settlement was always retained in the model to control for underling spatial variation in risk. Results are expressed as odds of having had Guinea worm and as relative risks, with 95% confidence intervals from bootstrapping 10,000 times with replacement. The same analyses were conducted on our field survey records of infection and the longer-term data collected by the CGWEP. For the models of CGWEP data, some (<5%) bootstrapped models did not converge and were excluded from calculations of confidence intervals.

Model selection adopted an information theoretic approach. A difference in Akaike Information Criterion (ΔAIC) of <2 was used to select the top model set. Unless otherwise stated, full model-averaged effect size coefficients with 95% confidence intervals and p-values from full model-averaged tables are reported. Correlations between explanatory variables were investigated prior to analyses and correlated variables were precluded from appearing in the same models. Analyses were undertaken in R (v.3.3.3) and Quantum Geographic Information System (v2.18.1). Locations were projected into the EPSG (32634) coordinate system. lme4 (v1.1-12) [16] was used to conduct GLMMs, MuMIn (v1.15.6) [17] for model selection and mgcv (v1.8.12) [18] for GAMs.

## Results

### Dog tracking

150 dogs were collared and 134 generated usable tracking data (Tables 1–2). Mean deployment was 7.7 days (SE=0.2). Dog ranging was significantly affected by body condition, settlement and the number of days for which a dog was monitored (Table 3); age and sex appeared in the top model sets but did not contribute significantly to variation after model averaging. Dogs in poor body condition had smaller total ranges (Mean = 10.50 km^2^ ± SE Mean 2.29 km^2^) than those in moderate condition (11.74 ± 1.91 km^2^; Effect size for logged range (95 % CIs) = −0.45 (−0.89, 0.00), p<0.05). The core and total ranges of dogs in Largana and Magrao were significantly smaller than those of dogs living in Kakale (Table 2; Largana vs Kakale: Effect size for logged core range: −0.89 (−1.69, −0.10), p<0.05; total range: −1.52 (−2.08, −0.95), p<0.001; Magrao vs Kakale: Effect size for logged core range: −2.00 (−2.71, −1.30), p<0.001; Effect size for logged total range: −1.47 (−1.97, −0.96), p<0.001). Use in hunting activity and household provision of water did not appear in any of the top models for variation in range areas (Table 3).

**Table 1.** Summary of dog characteristics in three settlements in Chad. Data are from all collared dogs (n=150), but some dogs have missing data for individual characteristics. Water provision is the proportion of households which reported providing water for animals.

| Settlement | n dogs | Sex<br>(M:F) | Body condition<br>(Poor:Moderate) | Mean age<br>(months) | Age range<br>(months) | Water provision<br>(%) |
| --- | --- | --- | --- | --- | --- | --- |
| Largana | 34 | 16 : 18 | 22 : 12 | 32 ± 4 | 6 - 96 | 60 |
| Kakale | 52 | 26 : 26 | 25 : 27 | 35 ± 3 | 4 - 144 | 75 |
| Magrao | 64 | 40 : 24 | 16 : 42 | 34 ± 3 | 5 - 144 | 100 |
| All | 150 | 82 : 68 | 63 : 81 | 34 ± 2 | 4 - 144 | 82 |

**Table 2.** Summary of dog ranging behaviour in three settlements in Chad. Core range is the 60% kernel density estimate and total range is the 100% minimum convex polygon. The mean and standard error in parentheses are presented for each variable.

| Settlement | n dogs | Core range<br>(km<sup>2</sup>) | Total range<br>(km<sup>2</sup>) | Fixes<br>within<br>100m of a<br>pond<br>(%) | Unique<br>ponds<br>visited | Ponds in<br>core<br>range | Ponds in<br>total<br>range | Fixes within<br>100m of the<br>Chari River<br>(%) |
| --- | --- | --- | --- | --- | --- | --- | --- | --- |
| Largana | 31 | 0.30(0.09) | 4.70(1.33) | 11(4) | 2(0.3) | 2(0.3) | 4(0.8) | 4(2) |
| Kakale | 48 | 2.53(0.74) | 21.01(3.08) | 36(5) | 8(0.6) | 8(1.3) | 25(3.1) | <1(<1) |
| Magrao | 55 | 0.17(0.04) | 4.39(0.63) | 18(4) | 3(0.3) | 1(0.2) | 6(0.7) | 2(1) |
| All | 134 | 1.05(0.28) | 10.42(1.35) | 23(3) | 4(0.3) | 3(0.6) | 12(1.4) | 2(<1) |

**Table 3.** Summaries of analyses of variation in dog range sizes, time spent by dogs in the vicinity of ponds and history of Guinea worm infection in dogs in rural Chad. Core range is the 60% kernel density estimate and total range is the 100% minimum convex polygon. History of Guinea worm infection is from our field survey records and records held by the Chad Guinea Worm Eradication Program. Model outcomes are shown as the top model sets (ΔAICc < 2 from the top model) for all analyses.

| Formula | df | LogLik | AICc | $\Delta AICc$ | Weight |
| --- | --- | --- | --- | --- | --- |
| **Total range size (100 % MCP)** |  |  |  |  |  |
| Days monitored + body condition + settlement | 6 | -165.34 | 343.5 | 0 | 0.54 |
| Days monitored + body condition + settlement + age | 7 | -165.01 | 345.1 | 1.62 | 0.24 |
| Days monitored + body condition + settlement + sex | 7 | -165.06 | 345.2 | 1.71 | 0.23 |
| **Core range size (60 % KDE)** |  |  |  |  |  |
| Settlement | 4 | -204.48 | 417.3 | 0 | 0.33 |
| Settlement + days monitored | 5 | -203.80 | 418.2 | 0.85 | 0.22 |
| Settlement + body condition | 5 | -203.91 | 418.4 | 1.07 | 0.19 |
| Settlement + days monitored + body condition | 6 | -203.17 | 419.1 | 1.81 | 0.13 |
| Settlement + water provision | 5 | -204.31 | 419.2 | 1.85 | 0.13 |
| **Time spent in the vicinity of ponds (n fixes within 100 m of ponds)** |  |  |  |  |  |
| Body condition + settlement + water provision + age | 7 | -1357.02 | 2729.0 | 0 | 0.54 |
| Body condition + settlement + water provision | 6 | -1358.29 | 2729.3 | 0.29 | 0.46 |
| **Guinea worm infection (reported history of infection or not)** |  |  |  |  |  |
| *Our field records (main analysis)* |  |  |  |  |  |
| % fish consumed + water provision | 3 | -52.34 | 110.9 | 0 | 0.52 |
| % fish consumed + water provision + settlement | 4 | -52.18 | 112.8 | 1.85 | 0.21 |
| *Longer-term CGWEP records (for validation)* |  |  |  |  |  |
| Water provision + settlement | 3 | -49.89 | 106.0 | 0 | 0.54 |
| Water provision + settlement + % fish consumed | 4 | -49.70 | 107.8 | 1.78 | 0.22 |

Using satellite imagery from earlier in the dry season, 342 ponds were identified in the three search areas. 30 ponds were located using GPS in the field, of which 17 were not detected using satellite imagery, suggesting that new ponds were forming as the rains commenced. A sample of 359 ponds was used in our analyses. On average, dogs had 23% of their locations within 100 m of a detected pond and 2% of their relocation points within 100 m of the Chari River. Dogs in poor body condition spent more time in the vicinity of ponds than those in moderate condition (Effect size in negative binomial model = 0.72 (0.28, 1.15), p<0.01; n=110). Water provision for animals was associated with dogs spending less time in the vicinity of ponds (−1.01 (−1.61, −0.42), p<0.001). Dogs from Kakale and Magrao spent more time in the vicinity of ponds than dogs from Largana. In Magrao, 37 of 101 (37%) ponds were visited by at least one dog, while in Kakale, 87 of 202 (43%) ponds were visited. A small minority of ponds accounted for the great majority of dog activity. For example, in Kakale, 80% of dog visits occurred at nine (5%) ponds, all of which were <100 m from a household with a tracked dog and 95% of visits occurred at 17 (8%) ponds, 16 of which were <200 m from a household (Figure 2).

**Figure 2.**
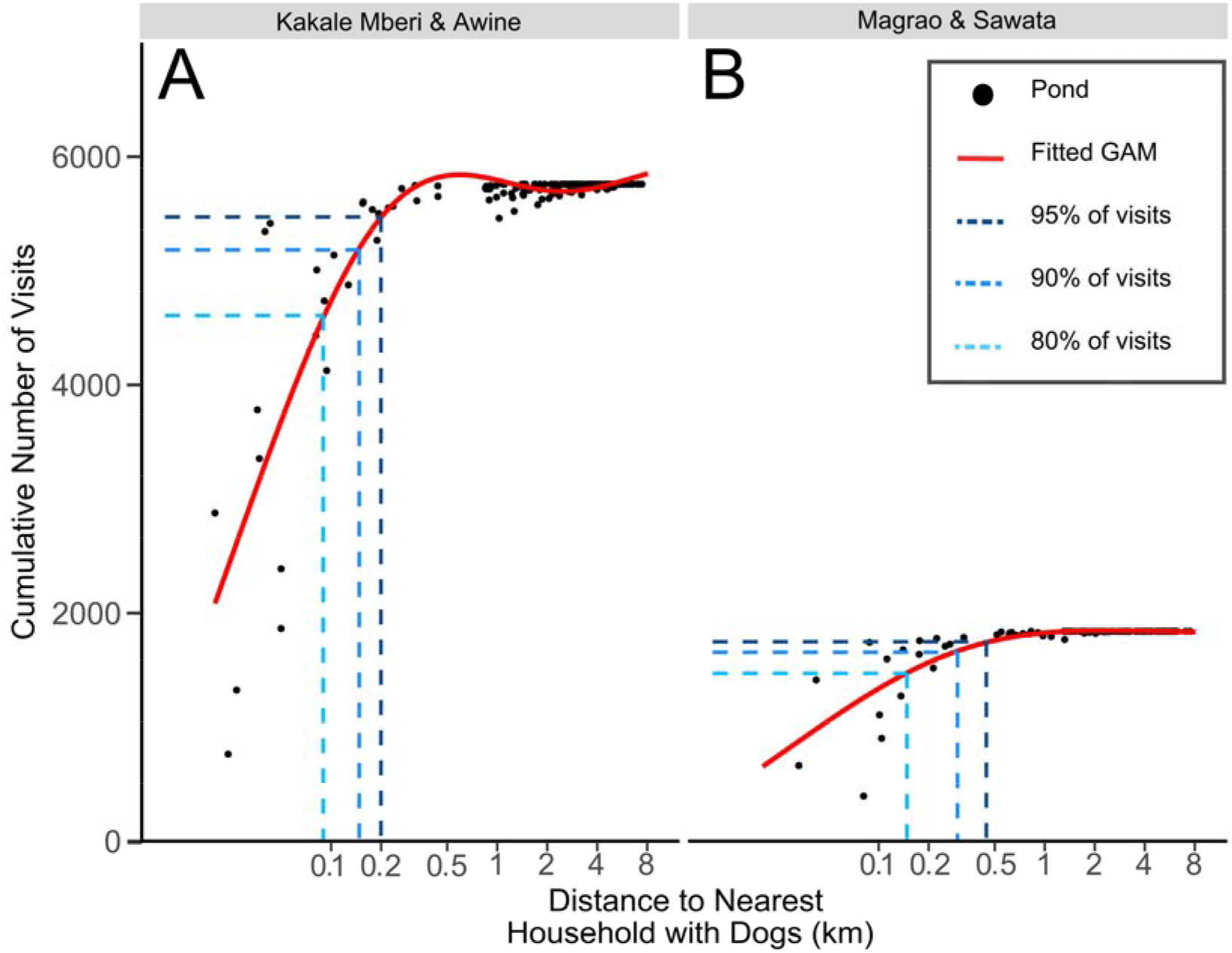
Distances from human households of the ponds most exposed to visits by dogs in two settlements in Chad. A is Kakale and B is Magrao. Dog visits were determined by locating dogs within 100m of a pond by GPS tracking. Ponds were located from satellite imagery within a search area enclosing all dog location points for each village and are a sub-sample of all available ponds. All visits by all dogs were summed for each pond and the cumulative total of all dog visits is plotted against the distance of the pond to the nearest household with tracked dogs, with the red line from a fitted generalised additive model. The dotted lines indicate the distance from a household with dogs at which 80%, 90% and 95% of all dog visits are captured.

### Dog diet

From 92 households, we recorded 346 reports of food items, grouped into seven broad, isotopically-distinct categories for which we collected 174 samples: C3 plant food (e.g. potatoes, peanuts, rice, n=9), C4 plant food (e.g. millet, sorghum, maize, n=17), livestock meat (n=25, including samples of 7 cows, 7 sheep, 9 goats, 2 chickens), wild animal meat (n=24, including samples of 17 amphibians, 2 reptiles, 2 mammals, 3 birds), fish (n=88) and human faeces (n=11). Whiskers were obtained from 108 dogs from Kakale and Magrao. Mean whisker length was 50.6 mm (SD 9.1), mean growth rate was 0.42 mm/day (SD 0.21, n=21), hence the period represented by the average whisker was ~120 days. C4 plant food (“boule”, a sorghum or millet porridge), human faeces and C3 plant food comprised the majority of dog diet and other items were eaten relatively infrequently, but in similar measure (Figure 3).

**Figure 3.**
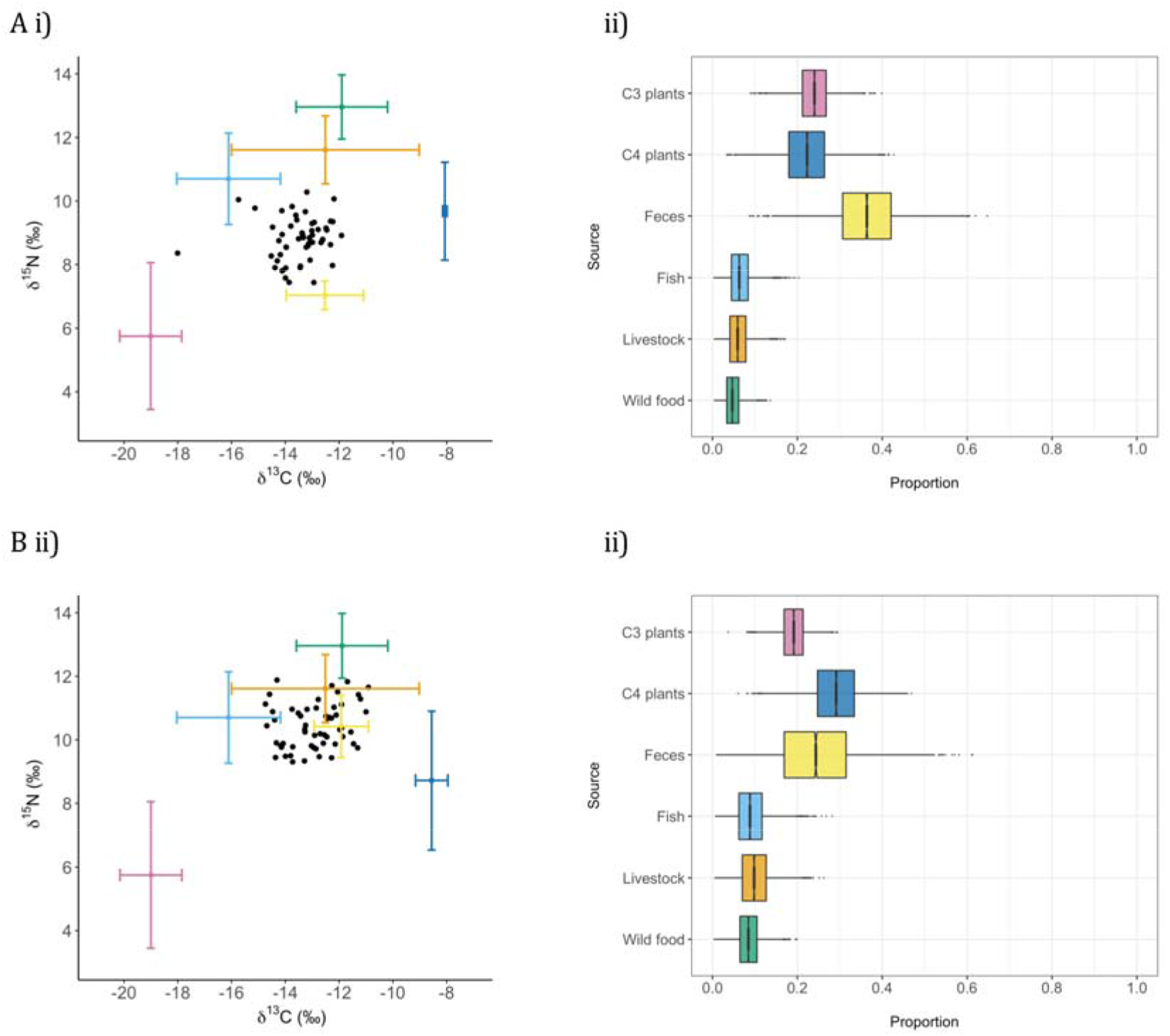
Stable isotope analysis of the diets of dogs in two settlements in Chad. Settlements are A) Kakale and B) Magrao. i) The δ^15^N and δ^13^C values for sampled dogs (black dots) and the mean ± standard deviation of δ^15^N and δ^13^C for putative food groups. Trophic discrimination factors, derived from package SIDER, have been applied to adjust dog isotope ratios downwards for both δ^15^N and δ^13^C. ii) Outputs of Bayesian mixing models, from package SIMMR, of the proportional contributions of the main food groups to dog diets.

### Guinea worm infection

Infection data were collected for 149 dogs, of which 34 (23%) had a history of Guinea worm emergence. A model including all explanatory variables was initially run (n=86), however body condition, age, sex or any spatial variables appeared in fewer than 50% of the top models and so dogs with missing data for these variables were included in the final model. Using our field data, 13 of 51 (25.5%) dogs in Magrao and 14 of 49 (28.6%) in Kakale had some history of Guinea worm infection. Dogs in households where water was provided for animals were less likely to have had Guinea worm (Figure 4; Table 3; Relative risk = 0.33 (0.11, 0.63) p<0.01), whereas dogs with a higher proportion of fish in their diets were more likely to have had Guinea worm (Figure 4; Table 3; Relative risk = 1.14 (1.00, 1·48), p < 0.05; Figure 4). CGWEP surveillance data recorded that 15 of 52 (28.8% dogs in Magrao and 8 of 50 (16.0%) dogs in Kakale had some history of worm infection. Analyses using CGWEP data suggested an effect of settlement (Increased risk in Magrao, compared to Kakale, Relative risk = 2.17 (0.19, 11.44), a similar effect of water provision (Relative risk = 0.40 (0.06, 0.71) but no effect of fish consumption (Tables S1-3).

**Figure 4.**
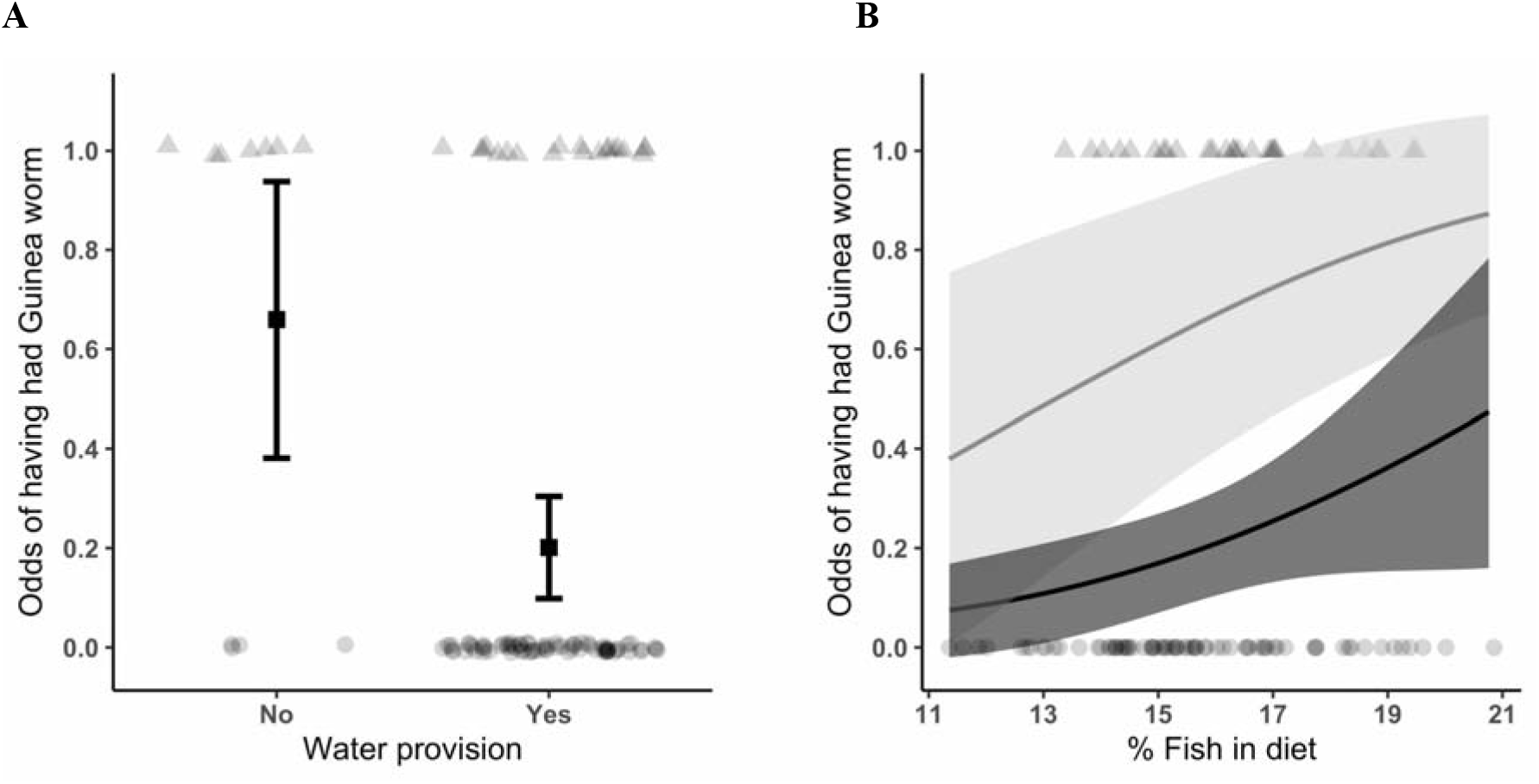
Effects of water provision and of fish consumption on Guinea worm infection history in dogs in Chad. The odds, with 95% confidence intervals, of having had Guinea worm are shown for dogs A) living in households in which animals are, and are not, provided with water, and B) with different modelled proportions of fish in their diets. The effects of fish consumption are presented separately for dogs living in households, in which animals are (dark grey) and are not (light grey) provided with water. Odds are from exponentiated model coefficients describing the relationship between the history of Guinea worm infection from our field survey, averaged across the top model set and for the different levels of values of predictive factors. Results presented are from dogs in Kakale but dogs in Magrao show the same relationships.

The median number of emergent worms per infected dog was 2. Two dogs with exceptionally high worm burdens (9 and 14 worms) had particular influence on analyses and when they were excluded, none of the explanatory variables had an effect on the number of worms that had emerged from an infected dog.

## Discussion

Human activity is a key influence upon domestic dog ecology in these settlements and relates to variation in ranging, diet, access to water and, critically, history of Guinea worm infection. Dogs spent most of their time close to their human households and their core ranges tended to be less than 1 km^2^. The exception was for dogs in Kakale, whose owners were moving frequently between permanent and seasonal residences for crop cultivation.

Dog exposure to ponds and their infection history were functions of the provision of water to animals by the household. We have shown that household provision of water to animals was protective of Guinea worm infection, and that water provision was negatively related to time spent close to ponds. There is a question of cause and effect here; the availability of water within the household might reduce the tendency for dogs to visit ponds but it is also likely that households located further from standing water might more frequently provide water for their animals. We did not determine the source of water provided for animals and analysis of whether water comes from safe, borehole sources or unsafe surface water sources may help refine our understanding of risk. Clearly, however, our findings highlight the importance of variation in where and how dogs access water in determining their exposure to Guinea worm infection.

In terms of potential sources of infection, a small number of ponds close to households accounted for almost all exposure of dogs to surface water, while visits to the Chari River were comparatively rare. Dog exposure to ponds close to settlements means that such ponds merit targeted investigation and prioritisation for systematic treatment with temephos (Abate). Dogs acquiring infection by drinking water contaminated with infected copepods would fit with a classical route for Guinea worm transmission. The inconsistency of this transmission pathway with very low incidence in humans can be explained by the relative ease of access to borehole water. Our analyses of satellite imagery reveal problems in accessing contemporary imaging at an adequate resolution and in reliably identifying ponds using optical images. Therefore, we suggest that additional techniques, such as radar, aerial and ground-based surveys, may assist in understanding the distribution of surface water in and around villages badly affected by Guinea worm.

Dogs relied heavily on anthropogenic foods, principally boule and human faeces. Given the similarity in the stable isotope ratios of these items, there is uncertainty in the exact contribution they each make, though together they clearly make up the majority of dog diet [19]. Although we were successful in determining between-dog variation in fish consumption, we could not distinguish amphibians from other wild animal meats in these settlements. We had strong *a priori* interest in consumption of aquatic foods, as a potential route for exposure of dogs to infection. We identified a significant effect of fish consumption on Guinea worm infection. There was considerable uncertainty associated with this effect, though over the range of observed values the risk increased markedly. This may highlight the importance of novel routes for exposure of dogs to infection, perhaps by the consumption of fish, fish guts or other remains.

It is, of course, possible that dogs are exposed to Guinea worm infection by both classical and novel routes, or perhaps that novel routes explain occasional introductions and the classical route “amplifies” infection in non-human hosts, in a similar manner as was once seen with humans.

We were able to conduct exploratory epidemiological analyses using history of adult female Guinea worm emergence as an indicator of infection. The underlying assumption of these analyses is that contemporary investigations reflect the ecology of the dog at the time at which it acquired Guinea worm infection. This may not be unreasonable, given dog longevity and consistency of their environment. However, our study, undertaken at the beginning of the rainy season, missed the major environmental variation arising from rains and flooding. We also took emergent adult worms as the only measure of infection, which omits dogs that are truly infected but have not yet progressed to worm emergence (i.e. some animals classified as uninfected were genuinely infected) and dogs that have been exposed (i.e. exhibit the ecological traits of infected animals) but have not progressed to infection. An effective diagnostic of prepatent infection in dogs could potentially reduce this classification uncertainty.

It is perhaps remarkable that we have, with even this relatively small-scale study, identified correlates of Guinea worm infection in these new non-human animal hosts. We hope our work provides encouragement that the impressive progress towards dracunculiasis eradication might continue apace, by addressing this new but now demonstrably tractable epidemiological setting in which the Guinea worm has emerged.

## Acknowledgements

Financial and logistic support was provided by The Carter Center (cartercenter.org), Chad Ministry of Public Health (sante-tchad.org) and World Health Organization (who.int). The Bill and Melinda Gates Foundation (gatesfoundation.org) provided satellite imagery. With the exception of the named authors affiliated with the funding organisations, the funders had no role in study design, data collection and analysis, decision to publish, or preparation of the manuscript. We thank Stuart Bearhop, Io Blair-Freese, Chris Cleveland, Melinda Denson, Mark Eberhard, Donald Hopkins, Rich Inger, Aileen Mill, Ken Neal, Philip Tchindebet Ouakou, Sharon Roy, Ernesto Ruiz-Tiben, Jordan Schermerhorn, Adam Weiss, Michael Yabsley, Hubert Zirimwabagabo and colleagues at University of Exeter for their advice and support.

